# Detection of Selenium/Sulphur substitution in the heterologous expression of Thioredoxin isoform 1 (Trx1) from *Saccharomyces cerevisiae*

**DOI:** 10.1101/2021.11.05.467525

**Authors:** Humberto Antunes de Almeida Filho, Leticia Miranda Santos Lery, Ana Paula Valente, Luis Eduardo Soares Netto, Fabio Ceneviva Lacerda de Almeida

## Abstract

Thioredoxins are ubiquous proteins with 2 cysteines at the active site. The isoform 1 of Thioredoxin from *Saccharomyces cerevisae* (Trx1) has six sulphur aminoacids, two cysteines and four methionines. In this work we performed the replacement of cysteines by selenocysteines by growth of a transformed celular expression vector *E. coli BL21-DE3* in selenocysteine containing culture medium. The Maldi-TOF spectra of Seleno/Sulphur substituted Trx1 revealed six component peaks with 46-48 Da range between them, that is the isotopic Seleno-Sulfur difference, showing the replacement of the Cysteines and Methionines to Selenocysteines and Selenomethionines. The Maldi-TOF spectra of the peptides derived from Trypsin digestion of the purified Thioredoxin (peptide mass fingerprint) show Selenocysteine and Selenomethionine containing peptides. Therefore we are demonstrating that cystein can be replaced by selenocystein and be metabolically converted to selenomethionine during Trx1 heterologous translation. Furthermore, the Maldi-TOF spectra are showing the presence of the most abundant isotopes of selenium inserted in the peptides containing cysteine and methionine, derived from the Trx1 digestion. The one dimensional ^77^*Se*–^1^*H* heteronuclear multiple quantum coherence NMR spectroscopy (1D-HMQC) for reduced Seleno substituted Trx1 (Se-Trx1), revealed three ressonance lines for ^1^*H*_*β*1_ from Selenocysteines 30 and 33, between 1.6 and 2,0 ppm. The bidimensional HMQC spectra (2D-HMQC) of the reduced Se-Trx1 show the ^77^ Se ressonance signal in 178 ppm, coupled with ^1^*H*_*β*1_ and ^1^*H*_*β*2_ lines between 2.1 and 1.8 ppm. The 1D-HMQC for oxidized Trx1 revealed the only one broad resonance in 2.6 ppm probably relative to the ^1^*H*_*β*1_ prótons. The 2D-HMQC spectrum of oxidized protein shows a higher chemical shift of selenocysteine ^77^Se (832 ppm) if compared to reduced state (178 ppm). Together these data are showing that the protocol of *Se – S* substitution developed here is a efficient method to label the active site of Thioredoxin 1 with a broad band chemical shift atom ^77^*Se*. Furthermore the large spectral window of the ^77^*Se* NMR detected between reduced and oxidized states of the Thioredoxin 1 shows that this atom is an excellent probe for accessing oxidative states and probably the conformational dynamics of the active site of the Se-Trx1.

## Introduction

Thioredoxins are ubiquous low mass proteins (11-12 kDa) with conserved sequence in its redox active site (Trp-Cys-Gly-Pro-Cys). When thioredoxin is in reduced state, the active-site cysteines presents a dithiol group that is able to catalyze the reduction of disulfides in several cellular proteins. Thioredoxin was first identified as electron donor for ribonucleotide reductase an essential enzyme for DNA synthesis, in *Escherichia coli* [10]. The replacement of the two redox active cysteine residues in *Escherichia coli* Thioredoxin to selenocysteine enzyme (Se-Trx) was achieved by expression of the *E. coli* thioredoxin gene (trxA) in a cysteine auxotrophic *E. coli* strain in the presence of D,L selenocysteine [22]. Similarly biosynthetically replacing Met with SeMet is accomplished by culturing a Met auxotroph (DL41) of *E. coli* in a defined medium containing the selenium analog of Met, SeMet, as the sole source of Methionine [20,21]. The studies of selenium as a trace element in living systems has focused on selenium metabolism and in the isolation and characterization of a number of proteins that contain selenium in the form of selenocysteine ([21]). The genetic and translational machinery for inserting Selenocysteines in proteins has been characterized and it is observed that there are specific genes that code for expression of t-RNAs that accept L-serine and cotranslationally inserts selenocysteine in protein synthesis [11]. The metabolic machinery pathway of intrinsic *E. coli* selenoproteins has been characterized in detail, this pathway requires the products of four genes, selA, selB, selC, and selD. The selC gene product, for example, was identified as a selenocysteyl-tRNASec that cotranslationally delivers selenocysteine at the in-frame UGA codon [1,9]. The cluster of the genes from Selenoproteins biosynthesis compose a specific pathway to selenium. Likewise, Selenium can substitute sulfur through of unspecific selenium incorporation by conventional metabolic pathways [13]. Thus, the enzymes involved in the cysteine biosynthetic pathway can generate free selenocysteine. The cysteyl-tRNA synthetase can charge the tRNAcys with either cysteine or selenocysteine and it is inserted into proteins nonspecifically. Although selenocysteine is synthesized by *E. coli*, the free amino acid is not directly attached to the selC tRNA, but it can be esterified to tRNACys and then inserted randomly in many proteins in place of cysteine [27]. The biosynthesis of sulfur-containing amino acids provides an example of pathways which exhibit alternative means for various organisms to synthesize their own metabolites. Indeed, cysteine and homocysteine can be synthesized directly from reduced sulfur, or by the interconversion of these two metabolites. Homocysteine is then converted into methionine by a methionine synthase. The pathways to sulphur containing aminoacids possibly recognize analogous with Selenium and perform the enzymatic reactions.

An interesting issue derived of the substitution of the sulfur-containing aminoacids to selenium residues is that the ^77^*Se* isotope is sensitive to NMR spectroscopy. The ^77^*Se* has natural abundance of 7.6 percent, and spin I=1/2 in the nucleus, with a broad chemical shielding response [5]. Therefore it is highly sensitive to changes in the electronic environment, such as modification in bonding and conformation [19]. Consequently, ^77^*Se* NMR measurements, in combination with theoretical calculations of the magnetic shielding tensor, will provide new approaches to probe the role of the local environment in the reactivity of cysteines [8]. Furthermore,^77^Se NMR could be used to probe the enzymes active site and catalytic reaction mechanism that involves cysteines. Additionally the dynamic of the active site could be investigated at the same time. In this work we developed a efficient protocol to replace sulphur by Selenium in the sulphur aminoacids, through heterologous expression of the *Saccharomyces cerevisae* Trx1 in a *E. coli* expression vector. We found a high yield in expression of transformed Se-Trx1. The isotopic composition of the transformed proteins was characterized by Maldi-TOF mass spectrometry and we found the Selenium most abundant isotopes in cysteine and methionine modified residues. Furthermore, we perfomed heteronuclear ^77^*Se* –^1^*H* NMR spectroscopy experiments with modified Trx1 using natural Se isotopic composition, in oxidant and reduced environments. We find that ^77^*Se* has a large chemical shift between oxidated and reduced states of modified Se-Trx1. We assigned some heteronuclear NMR signals in one and two dimensions to ^1^*H*^*β*1^ and ^1^*H*^*β*2^ of Selenocysteines in the Se-Trx1. The large chemical shifts found between the oxidized and reduced forms of the modified protein is possibly generated by the diselenol-diselenide state of the selenocysteines of the active site of the modified Trx1. Therefore our results show that the S-Se substitution protocol developed in this work is a powerful tool for investigating oxidation states in enzymes that contain cysteines at the active site, through ^77^*Se* NMR spectroscopy. In addition, the high performance of the expression of modified proteins shows that our protocol can be applied in obtaining significant amounts of proteins for structural studies by different spectroscopic techniques.

## Materials and Methods

### The *Trx*1 Selenocystein labelling, expression and purification

The *E coli BL21-DE3* strain harboring the pET-17B/Trx1 and pLysE plasmids [23,24] stored at –80°c, were seeded in a petri dish containing rich culture medium, 32g/l LB-agar and cultivated at 37°c for 12h. Afterwards, a colony was isolated from the plate and inoculated in 2ml of LB medium (rich culture medium, without addition of agar) and cultured at 37°c until an optical density (*OD*_600_) of 1.0. This culture was used as an inoculum for 50 ml of M9 minimum culture medium and incubated overnight for 12 hours under agitation at 200 rpm. This culture was used as a next inoculum for 1l of M9. The D,L-cystine dipeptide (Sigma-Aldrich) 4.1 mM was reduced with Dithiotreitol (DTT) to 5 mM in a separate flask containing 50 ml of phosphate buffer pH = 7.2 and heated at a water bath in 90°c in order to obtain Selenocystein. This solution was added to 1l of the M9 culture medium as described previously. The ampicillin 100μg/ml was used in all culture media as a selection marker. The *E. coli* BL21DE3 cells obtained in the 50 ml of the M9 inoculum, were added to 1l of the M9 Selenocystein added medium described previously. The cells growth in 37°c under 200 rpm stirring until *OD*_600_ 1.0 during 4 hours, and 1 mM of Isopropyl β-d-1-thiogalactopyranoside (IPTG) was added in order to induce the protein expression. The protein expression was induced by 18 hours. The 1l of M9 containing cells described previously was centrifuged at 10, 000 × *g*. The cell pellet was collected and resuspended in 50 ml of lysis buffer containing 2mM *NaH*_2_*PO*_4_, pH=7.0, 3mM beta mercaptoethanol, 5μg/ml deoxyribonuclease, protease inhibitor cocktail (Sigma) (P2714), 16μg/ml lysozyme was then added and incubated for 30 minutes. Cells were then exposed to 6 cycles of 10 seconds of sonication at medium potency. The entire procedure was done at 0°c. The lysate was centrifuged at 4500 × *g* for 15 minutes and the supernatant was collected and heated to 90°c, and centrifugued again in 4500 × *g* for 10 minutes. The Trx1 refolds to a stable structure, after cooling [14], so the supernatant of the last centrifugation was collected. This material was applied to an ion exchange chromatography column containing DEAE-Toyo Pearl 650M resin (Toso Biosciences) previously equilibrated with 20mM Tris-HCl buffer pH=8.0, 3mM beta mercaptoethanol, 1mM sodium azide. Then a discontinuous elution gradient was made with NaCl at concentrations of 50 mM, 100 mM, 200 mM and 1 M in 20 mM Tris-HCl pH=8.0 3 mM beta mercaptoethanol, 1 mM sodium azide. The peak eluted in 50 mM NaCl was collected and dialyzed against 2 l of milli-Q water in 2 changes with an interval of 12 hours, and then lyophilized. This sample was subjected to analysis by SDS-PAGE where Trx1 was observed to be significantly preponderant although not completely purified. This fraction was then concentrated by ultrafiltration in amicom with a 3 KDa exclusion membrane, to a final volume of approximately 5ml. This sample was submitted to a new liquid chromatography step in a Shimadzu binary HPLC system through a size exclusion column (Superdex HR75 Pharmacia) using TRIS-HCl buffer 20mM NaCl 0.15M beta mercaptoethanol 3mM, *NaN*_3_ 1mM, as mobile phase in a single-phase solvent system with a flow of 1ml/min. The sample was applied to the column in volumes of 250μl in a time of 25 minutes for each chromatographic run. Protein elution was analyzed by absorbance detector (SPD-10AV Shimadzu UV-Visible). A preponderant peak corresponding to Trx1 was observed at a retention time of approximately 17 minutes. The fractions corresponding to this peak were collected in each step of the chromatographic run and grouped in a single sample. An aliquot of this sample was subjected to analysis by SDS-PAGE that showed a completely pure band with approximate mass of 12 KDa.

### Mass spectrometry

The MALDI-TOF mass spectra were obtained using a Voyager DE-PRO MALDI-TOF mass spectrometer (Applied Biosystems), operated in linear mode. Samples with 1*μl* of Se-Trx1 and Trx1 10μM were prepared on a stainless steel plate by the dried-droplet method using a matrix composed by Synapinic acid 10 mg/ml, 30% of *CH*_3_*CN* and 0.1 % of TFA. The mass calibration was performed using calmix 3 calibrant mixture (Applied Biosystems). The analysis of the mass pattern from peptides obtained by tryptic digestion of Trx1 (“Peptide Mass Fingerprint”) was made from the extract enriched with the protein, obtained after an expression protocol using Selenocystein in culture medium, as well as through the extract obtained by the protein expression without using Selenocysteine. The Trx1 and Se-Trx1 proteins were extracted from a SDS-PAGE gel containing pellet lysates of culture medium submited to electrophoresis. The band corresponding to the Trx1 protein was sliced and digested in gel by the enzyme trypsin. (Promega Madison, WI, USA), [17]. The theoretical mass pattern generated by digestion of Trx1 was performed by peptidemass software [25] in order to compare the effectiveness of the trypsin digestion protocol and the mass pattern generated by Maldi-TOF; The sample was mixed with a matrix of α-cyano 4 hydroxycinnamic acid in a solution of 50 % acetonitrile, 1 % trifluoroacetic acid, and analyzed in the Voyager DE PRO Biospectrometry Worksation (Applied Biosystems). The spectra were acquired in reflected mode with the device “delayed extraction”, in the mass/charge range of 800 to 4000 daltons; and analyzed in the data explorer v. program. 5.1. The internal calibrations were performed using tryptic peaks 2211.1046, 1045.5642 and 842.5 as reference and external calibrations using as standard a commercial mixture of peptides of known mass (Calmix1, Applied Biosystem). The background line corrections, noise filtering, selection of monoisotopic peaks and listing of well-resolved peaks were performed. The accuracy of the masses obtained in the spectra considered acceptable were better than 0.001 daltons (Da). Searches in databases were made through the MS-FIT program interface of the protein prospector [2].

### NMR samples preparing

#### Reduced Se-Trx1

The Se-Trx1 sample purified by HPLC as described above, was concentrated in an amicon ultrafiltration system (membrane with 3 kDa exclusion) to a final volume of 5 ml and subsequently in a centricon filter with the same exclusion size, through centrifugation at 4500 X g, for a final volume of 0.45ml, the procedure was repeated three times in order to change the buffer condition to 20mM *Na*_2_*HPO*_4_.7*H*_2_*O*, pH 7.2 added with 3mM deuterated dithiothreitol (*DTTD*), 3mM *NaN*_3_. 50*μl* of deuterated water (*D*_2_*O*). The concentration was estimated by spectrophotometry through the molar extinction coefficient in 6M of guanidine hydrochloride (GDNHCL) (*ϵtrx1* = 9530M^-1^) [7].

#### Oxidized Se-Trx1

The Trx1 sample prepared under reducing conditions as described above was oxidized with hydrogen peroxide (H_2_O_2_) at 5 mM, and had its buffer exchanged again to 20*mMNa*_2_*HPO*_4_.7*H*_2_O, pH = 7.2, through ultrafiltration in centricon. The sample was washed 10 times with the buffer in order to remove both excess *H_2_O_2_* and *DTT_D_* and 50*μl* of deuterated water (D_2_O) was added to the sample.

### NMR spectroscopy

#### ^1^*H* homonuclear NMR

One-dimensional ^1^*H* spectrum of the reduced Se-Trx1, were recorded at 303K temperature, 128 FIDs were collected with 4096 points in a spectral window of 9615.38 Hz.

#### ^77^*Se* –^1^*H* Heteronuclear NMR

Heteronuclear NMR spectra were collected on a Bruker DRX 600 Mhz in a double resonance probe tuned to ^77^*Se*. The HMQC spectra were made using a pulse sequence added with watergate [15]. We effected the signal modulation from ^77^*Se* NMR, by scanning the transmitter frequencies (O2P) to ^77^*Se*, in the heteronuclear resonance experiments (HMQC), using the signal of the dimethyl selenide *CH*_3_*SeCH*_3_ in deuterated chloroform (*CDCL*_3_), as the central reference of the spectral window. The frequency of the coupling constants used in the 1D-HMQC experiments ^2^*J*(^1^*H* –^77^ *Se*) were modulated according with the signal intensity and a coupling constant of the 13 Hz was adopted in the heteronuclear ^77^*Se* –^1^*H* HMQC experiments. It is in agreement with a ^2^*J*(^1^*H*_*β*1_ –^7 7^*Se*) constant coupling assigned in the 1,1 Selenocystine in previous works [18]. The signal intensity was observed in the spectra as a function of the coupling constants in order to observes the response o spin systems coupled with ^77^*Se*. The frequency of O2P was modulated in experiments with reduced and oxidized Se-Trx1 protein and the spectra were collected to optimize the intensities of coupled resonances. The detection was performed by quadrature and States-TPPI in the indirect dimension. The ^77^*Se* NMR signals were referenced to diphenylselenide compound at 463 ppm as a reference to dimethyl selenide compound in *CDCl*_3_.

The 2D-HMQC spectra ^77^Se edited, from reduced Se-Trx1 in 150μM concentration were collected at 303K temperature, using an O2P of 150 ppm, 6144 FIDs were collected in the direct dimension and 16 FIDs in the indirect dimension, with 1024 complex points in the direct dimension and 14 complex points in the indirect dimension (States-TPPI). The spectral window used was 9615.38Hz in the direct dimension and 11144 Hz in the indirect dimension. The spectra processing used the function of window with multiplication by cosine with zero padding up to 1024 points in the forward dimension and 64 points in the inverse dimension.

The 2D-HMQC spectra for oxidized Se-Trx1 in 570 concentration, were collected at 303K temperature using 02P at 800ppm. 3072 FIDs were collected in the forward dimension and 16 FIDs in the inverse dimension with 1024 points in the forward dimension and 14 complex points in the inverse dimension. The spectral window used was 9615.38Hz in the direct dimension and 17167 Hz in the indirect dimension. The spectra processing used the window function with multiplication by cosine with zero padding up to 1024 points in the direct dimension and 64 points in the inverse dimension.

## Results

### The Maldi-TOF characterization

The Trx1 computed mass based on aminoacid sequence is 11.234,98 Da (compute pI / Mw Tool [25]. Six sulfur containing aminoacids are found at sequence, two cysteines and four methionines (Figure 1 A). The mass measured to Trx1 expressed in Selenocystein suplemented culture medium by MALDI-TOF, show a complex mass pattern composed by 6 component peaks (fig 1B). The mass range between Maldi-TOF peaks revealed ranging between 45-48 Da, the same variation between selenium and sulphur atoms. Therefore the mass pattern fits with the substitution of sulfur by selenium in the six aminoacid residues of the Trx1.

**Figure 1.**
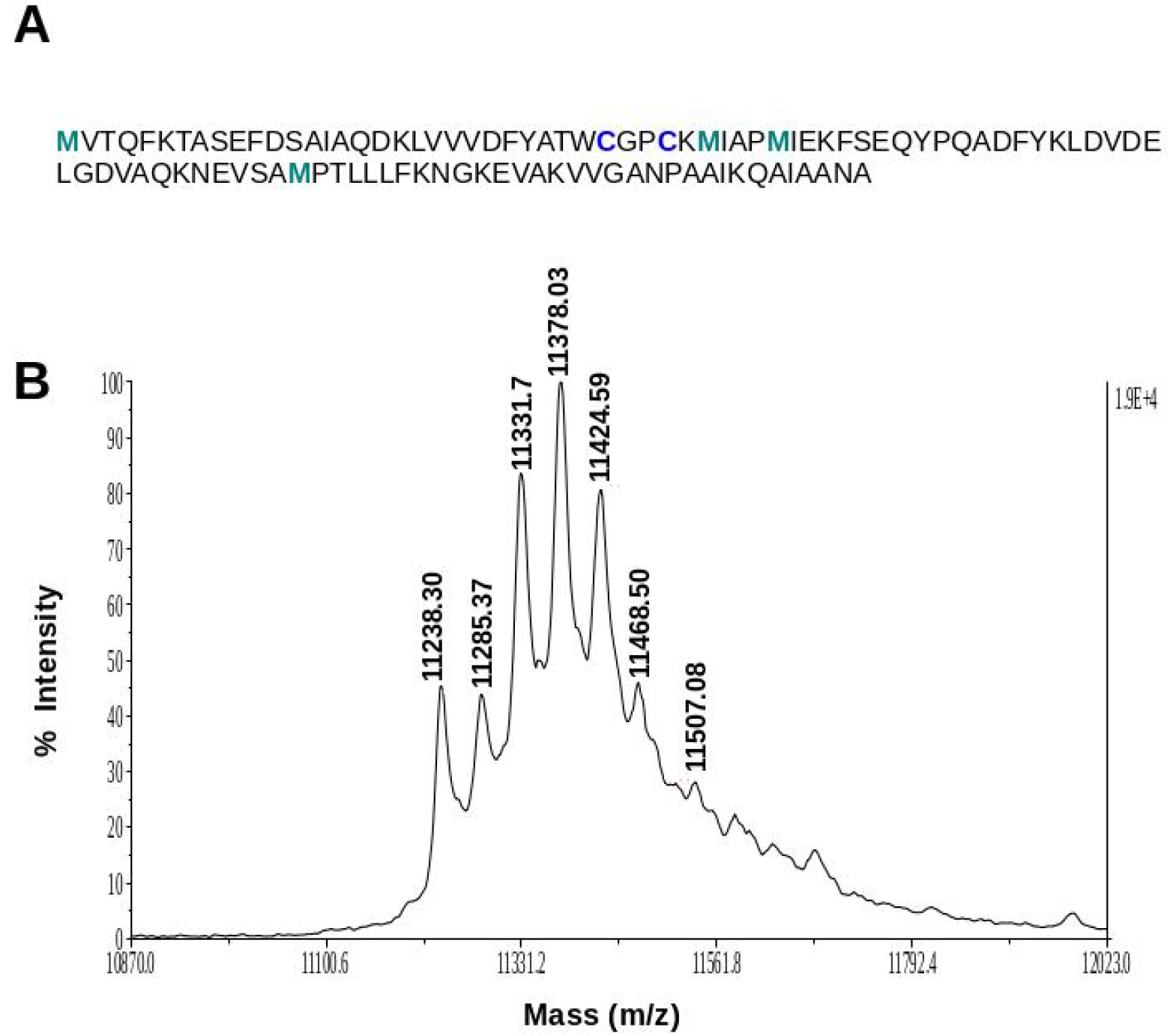
(A) Trx1 Aminoacid sequence indicating sulphur aminoacids (4 methionines (green) and two cysteines (blue). (B) MALDI-TOF peaks from biosubstituted Trx1 shown seven components peaks with 45-48 Da range corresponding to Trx1 (11238.30), 1Se-Trx1 (11285.37), 2Se-Trx1 (11331.7), 3Se-Trx1 (11378.03), 4Se-Trx1 (11424.05), 5Se-Trx1 (11468.5) and 6 Se-Trx1 (11507.08) revealing both selenocysteine and selenomethionine containing populations from Se-Trx1.

These results are in agreement with previous works that observes the cysteine-selenocysteine substitution in *E. coli* Thioredoxin (Trx) [12].

Previous works don’t reports the substitutions extensive to Methionines, therefore the detection of the selenocysteine to selenomethionine convertion by *E. coli* metabolic pathways are suggesting the steric recognition of enzymes such as cystathionine *β – synthase* and methionine synthase for metabolic intermediates containing selenium instead sulphur.

The Trx1 trypsin digestion was accomplished to confirm the biosubstitution by means of compare the mass of tryptic peptide fragments from biosusbtituted and non-biosubstituted Trx1. We performed a theoretical tryptic digestion from Trx1 generated with peptide mass software [25] (table 1), in order to compare the theoretical mass pattern against the one provided by Maldi-TOF. spectrometry (figure 2).

**Table 1.**
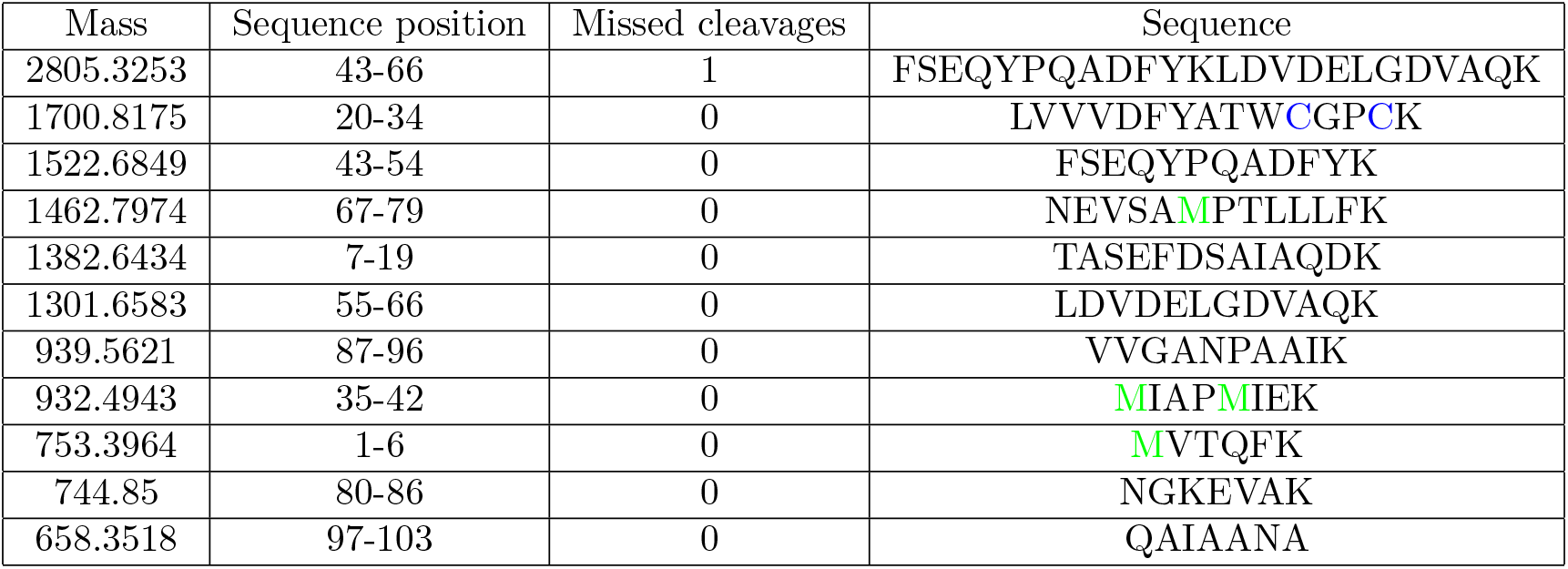
Peptides from Trx1 generated by theoretical Trypsin digestion with peptide mass software performing 0 and 1 missed cleavages

**Figure 2.**
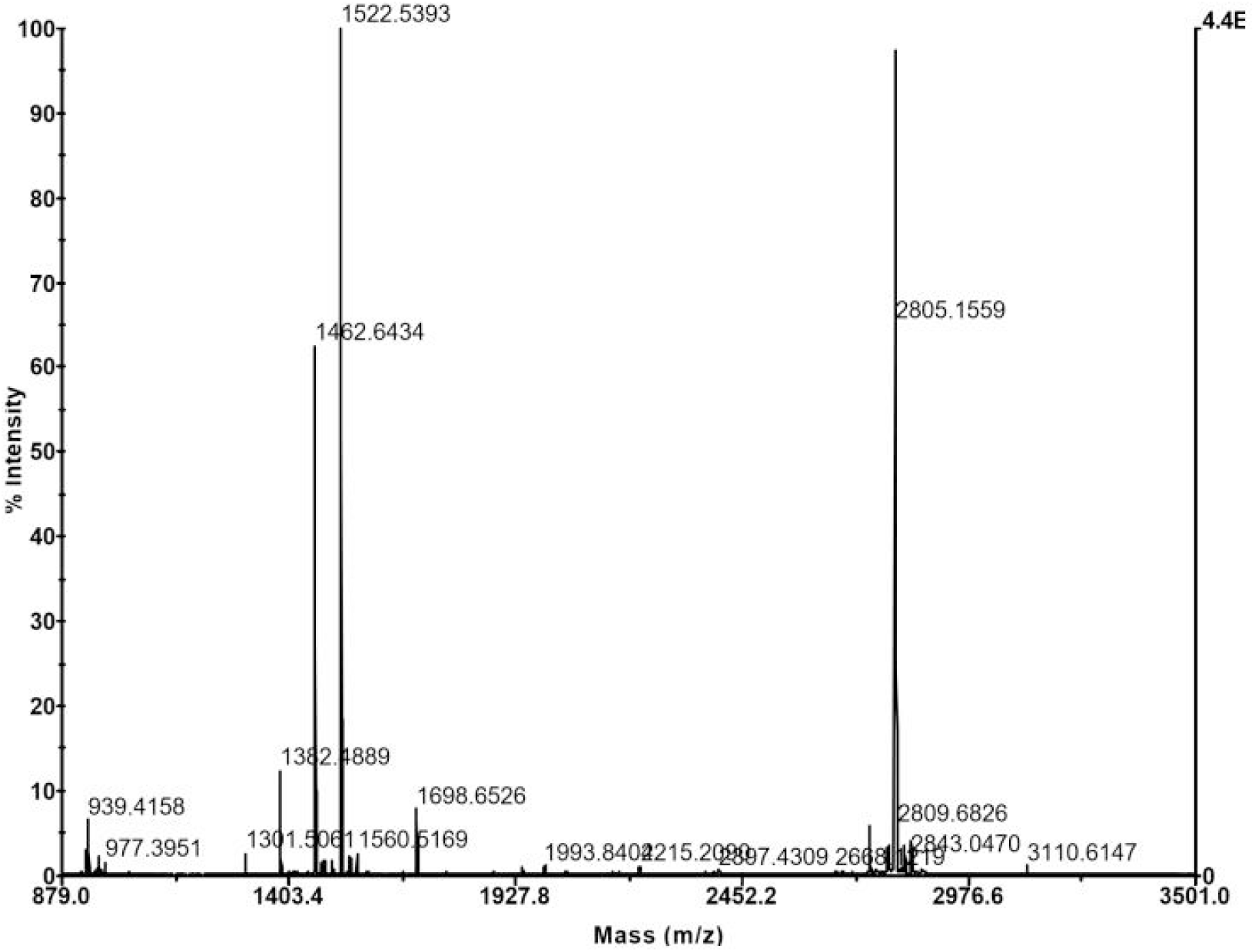
The Maldi-TOF Peptide mass fingerprint from tryptic digestion of Trx1

The peptide mass fingerprint obtained from Trx1 show a high agreement between theoretical and experimental mass of generated tryptic peptides. It confirms the effectiveness of the trypsin digestion protocol, so that the mass of peptides generated by tryptic digestion of Trx1 and Se-Tx1 can be compared to observe the substitution of sulfur by selenium with a greater precision.

The mass range to two Cystein containing tryptic peptide fragment (1698-1700 Da, see table 1 and fig 2) was probed in both Trx1 and Se-Trx1 (fig 3). The mass spectra to proteins show a complete substitution of two Cysteins by Selenocysteines with a addicional mass range of 96-98 Da to Se-Trx1, that is the same value to double Sulphur-Selenium replacement (fig 3 A), the maintenance of the original mass of this peptide in non substituted Trx1 is observed (fig 3 B).

**Figure 3.**
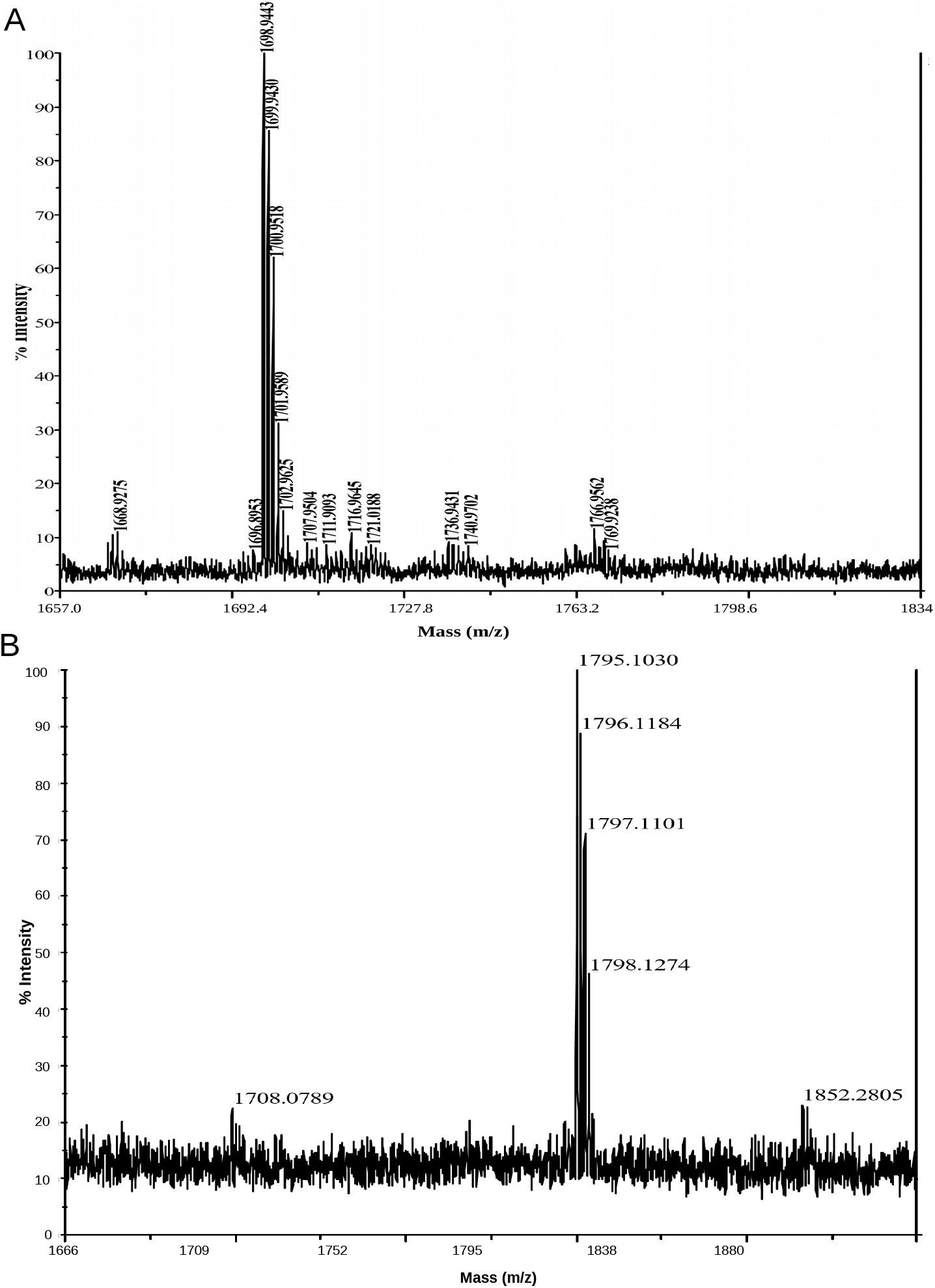
MALDI-TOF spectra from Two Cysteine containing peptide from Trx1 and Se-Trx1: (A) Spectra of two cystein containing peptide at non substituted Trx1 (B) Spectra from Se-Trx1 shown a average mass range between 94-98 Da between this peptide and the showed in A, It is corresponding to addition of two Se atoms at two cysteine containing peptide.

The high level of double cysteine/Se-cysteine substitution detected in the double cysteine tryptic peptide of the Se-Trx1 suggest that expression induction time of 22 hours in presence of M9 added Selenocystein (see methodology section) is able to generate a high level of biosubstitution.

The Maldi-TOF spectra from the methionine containing tryptic peptide at Se-Trx1 and Trx1 were compared (see table 1 and fig 4). The mass spectra revealed a cluster of peaks between 1506 and 1515 Da in the Se-Trx1 (see table 1 fig 4 B) that not appeared in Trx1 protein that just presented the peptide containing Methionine with approximately 1463 Da (See table 1 and fig 4 A). The cluster of peaks observed in the selenomethionine containing peptide is a clear evidence of the isotopic composition from selenium in the Se-Trx1, since Selenium has 6 isotopes with significative natural abundance: ^74^*Se*,^7 6^*Se*,^7 7^*Se*,^7 8^*Se*,^8 0^*Se* and ^82^*Se* [3]. The Subtraction between the the mass value of the most abundant isotopic peak 1511.03 showed in fig 4B and the peak referring to the mass of the peptide of 1463.08 Da, gives a value of 48 Da (*δ*M= 48) which is approximately equal to the subtraction of the masses of ^80^*Se* and ^32^*S*. It confirms the metabolic conversion between Selenocysteine and Selenomethionine during heterologous expression of Trx1 in a Selenocysteine containing M9. Together the MALDI-TOF data from Seleno/sulphur Trx1 show that different populations of Selenium labeled protein can be achieved in Selenocysteine added growth medium conditions.

**Figure 4.**
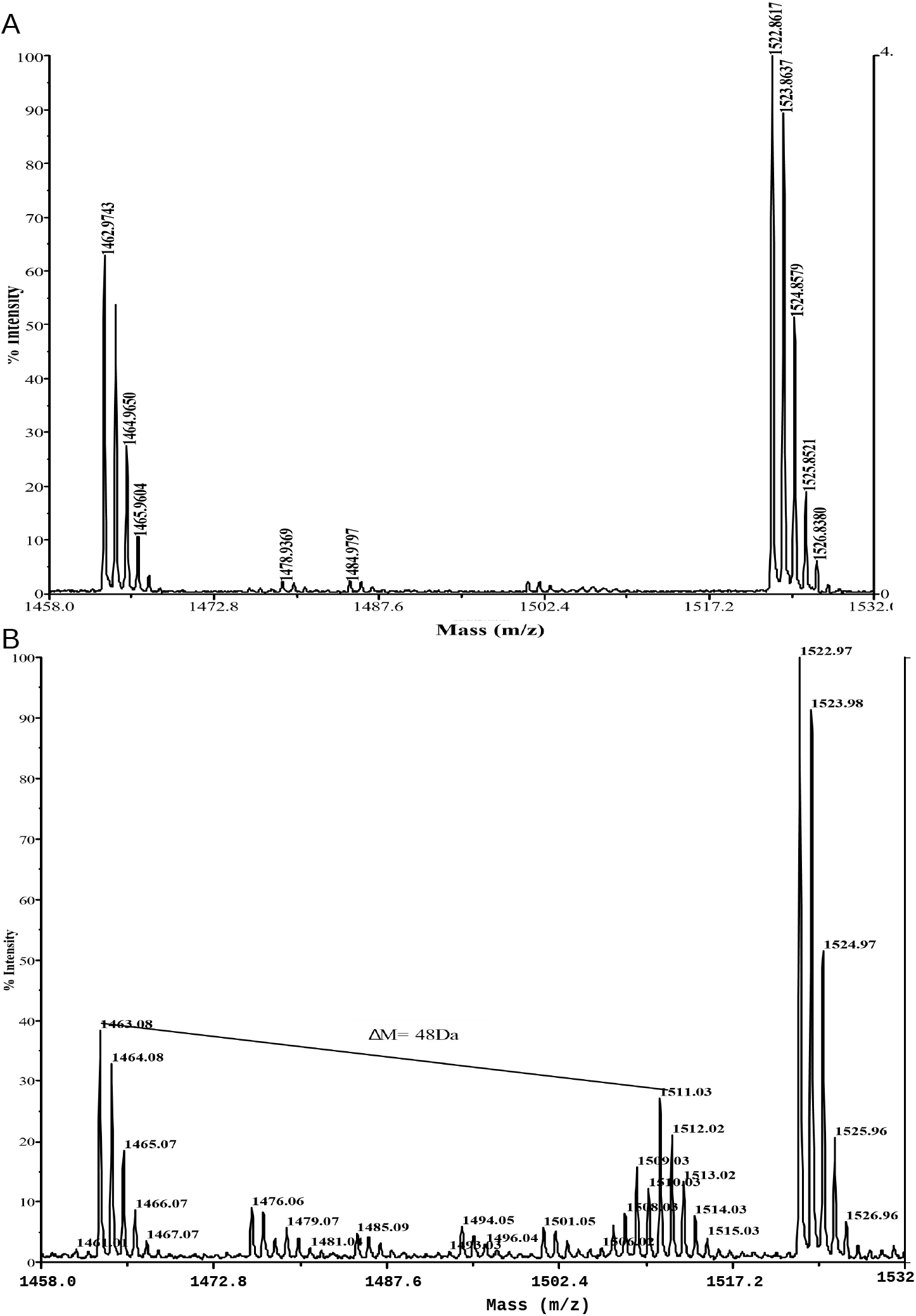
(A) The MALDI-TOF spectra of Trx1 tryptic fragments showing the mass of 1463 Da referring to the peptide containing a methionine in the primary sequence (See table 1). (B) The MALDI-TOF spectra of tryptic fragment of 1463 Da of the biosubstituted Se-Trx1 showing a additional set of peaks between 1506 and 1511 Da that were not present in the spectrum with the same mass range showed in (A).

### NMR spectroscopy characterization of the Se-Trx1

#### Reduced Se-Trx1

The one-dimensional ^1^*H* NMR spectra of the reduced Se-Trx1 reveals a pattern of well-resolved and dispersed resonances in the frequency range between 10 and 6 ppm which is an indication that the protein is folded and that the S-Se substitution did not unfolds the protein (See figure 5).

**Figure 5.**
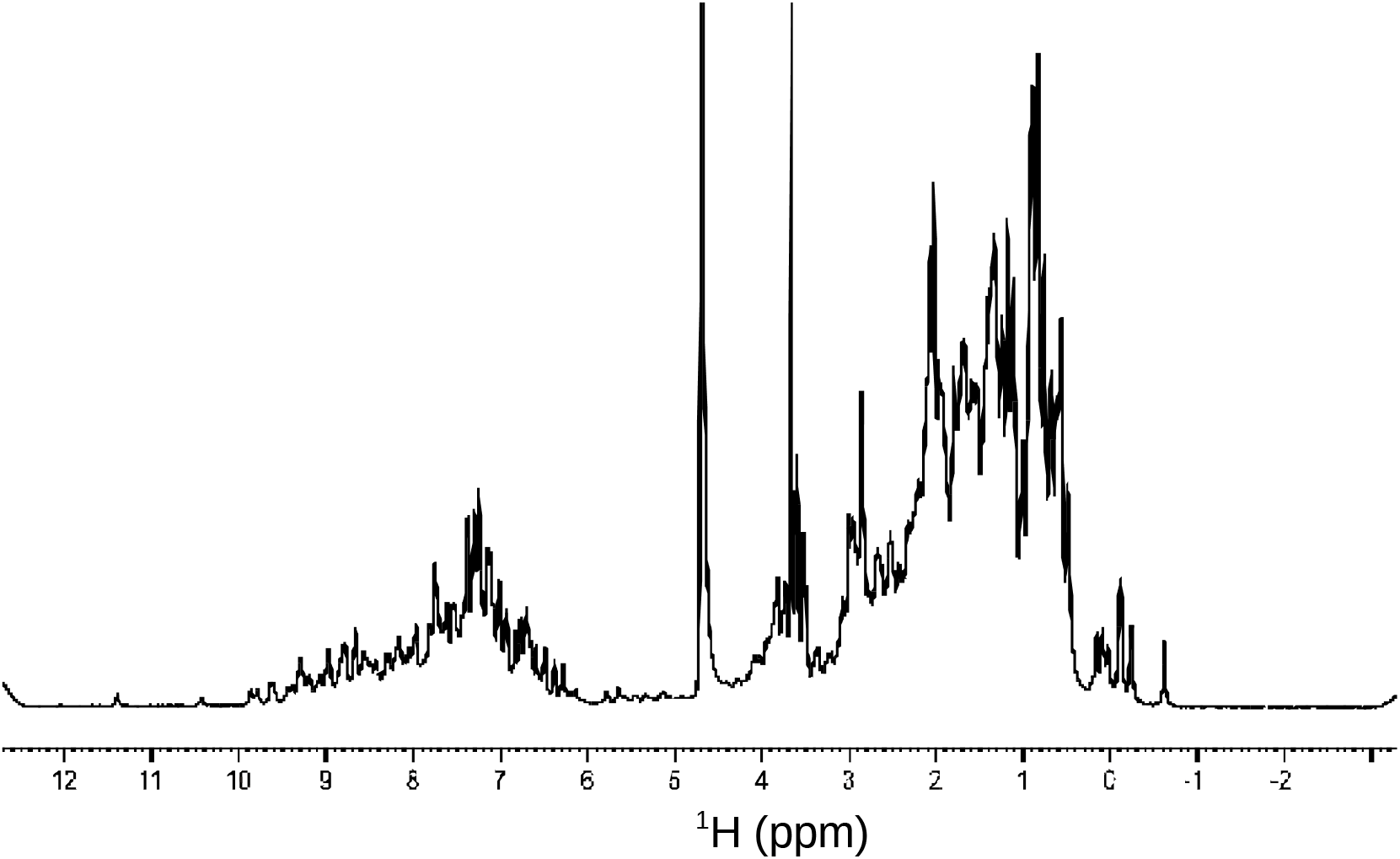
One-dimensional ^1^*H* spectrum of the reduced Se-Trx1 populations showing the resonant frequency of the amide hydrogens between 10 and 6 ppm, and a aliphatic hydrogens between 5 and 2 ppm.

We scan the ^77^*Se* signal in the HMQC heteronuclear experiments according with the frequency of the carrier wave excitation to ^77^*Se* (O2P) using a known (^2^*J* (*H*_*β*1_ 1 –^7 7^*Se*)) coupling constant to l,l-(^77^*Se*)_2_ cystine [18]. We tried to optimize the excitation the ^77^*Se* carrier wave (O2P) in order to obtain heteronuclear correlation spectra (^1^*H* –^77^*Se*) with better signal/noisy ratios. The resonance frequency for ^77^*Se* in clusters (*C* – *Se* – *H*) is between 200 and 500 ppm [4] and the resonance frequency to ^77^*Se* in selenocystin is 290 ppm [18]. Then we perform the HMQC experiments with O2P ranging between 100 and 300 ppm with a range of the 50ppm between experiments in order to observe the ideal excitation frequency inside this range. A coupled signal of the (^1^*H* –^7 7^*Se*) between 2 and 1.8 ppm was observed and it was strongly modulated by O2P (figure 6). The signals obtained with O2P at 150 and 200 ppm were more intense and this range were defined to the excitation of ^77^*Se*.

**Figure 6.**
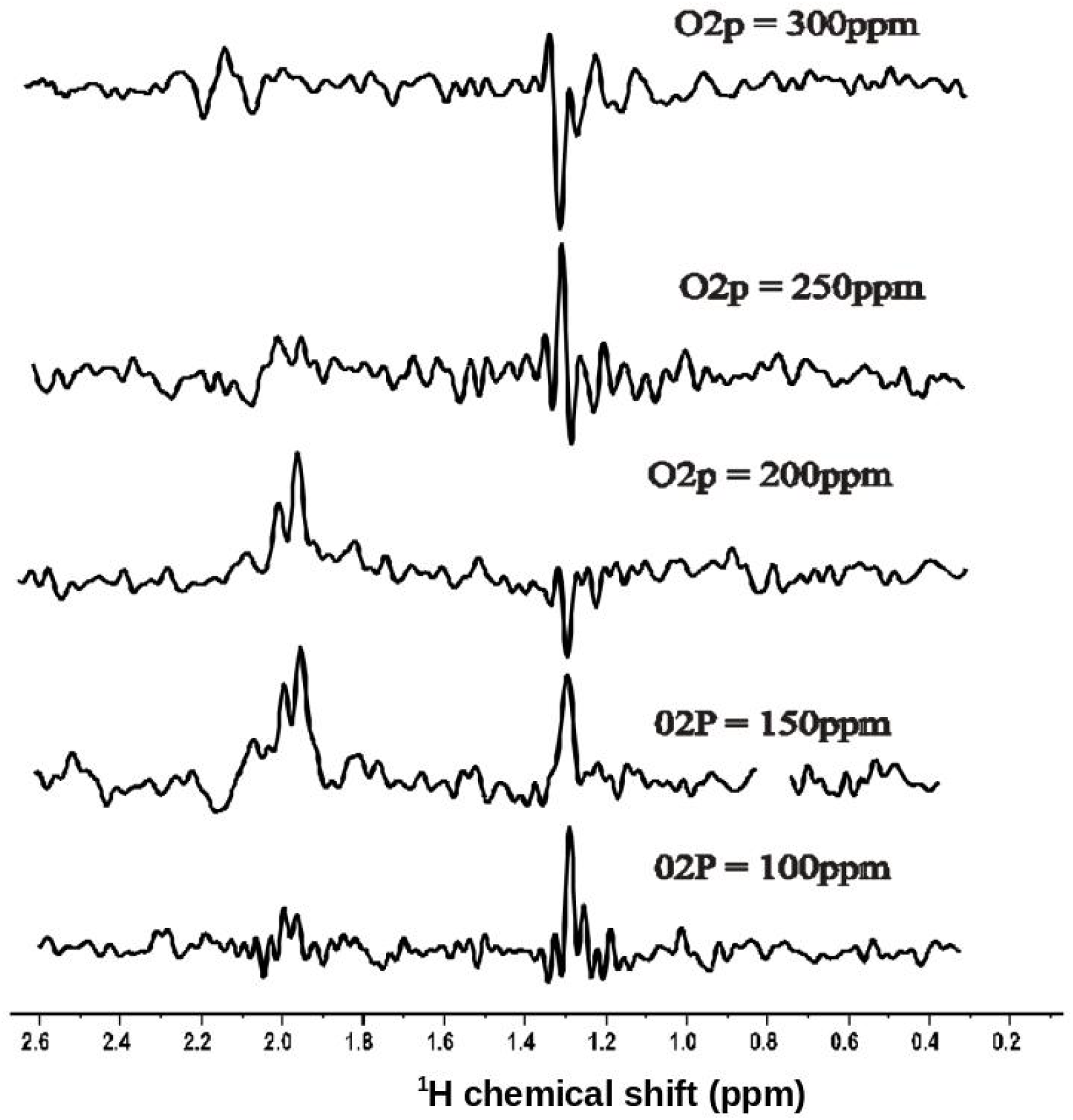
One-dimension HMQC spectra edited for ^77^*Se*. The spectra of 200 from reduced Se-Trx1 were collected with ^77^*Se* (O2P) excitation frequencies at 100ppm, 150ppm, 200ppm, 250ppm and 300ppm respectively. The spectra were collected at 303K with 7168 FIDs in a spectral window of 9615,384 Hz using a 13 Hz coupling constants.

The signal observed at 1.3 ppm is probably an artifact generated by phase errors in the HMQC. This artifact disappears when the FID accumulation is increased as evidenced in the following figures (figures 7 and 8). We performed the scanning in the coupling constants in order to assign the signal according with the coupling constant modulation. The resonance lines obtained between 2 and 1.8 ppm were observed as a function of the coupling constant. We observes better signal-to-noise ratios when using coupling constants between 10 and 16 Hz than between 18 and 25 Hz. (figure 7). This data strongly suggests that the resonance lines observed between 2 and 1.8 Hz refer to ^2^ J couplings to cystein ^1^*H*_*β*(1,2)_ since the coupling constant of the 13.1 Hz between this protons and ^77^*Se* is assigned in selenocystin, as already described in previous works. [18].

**Figure 7.**
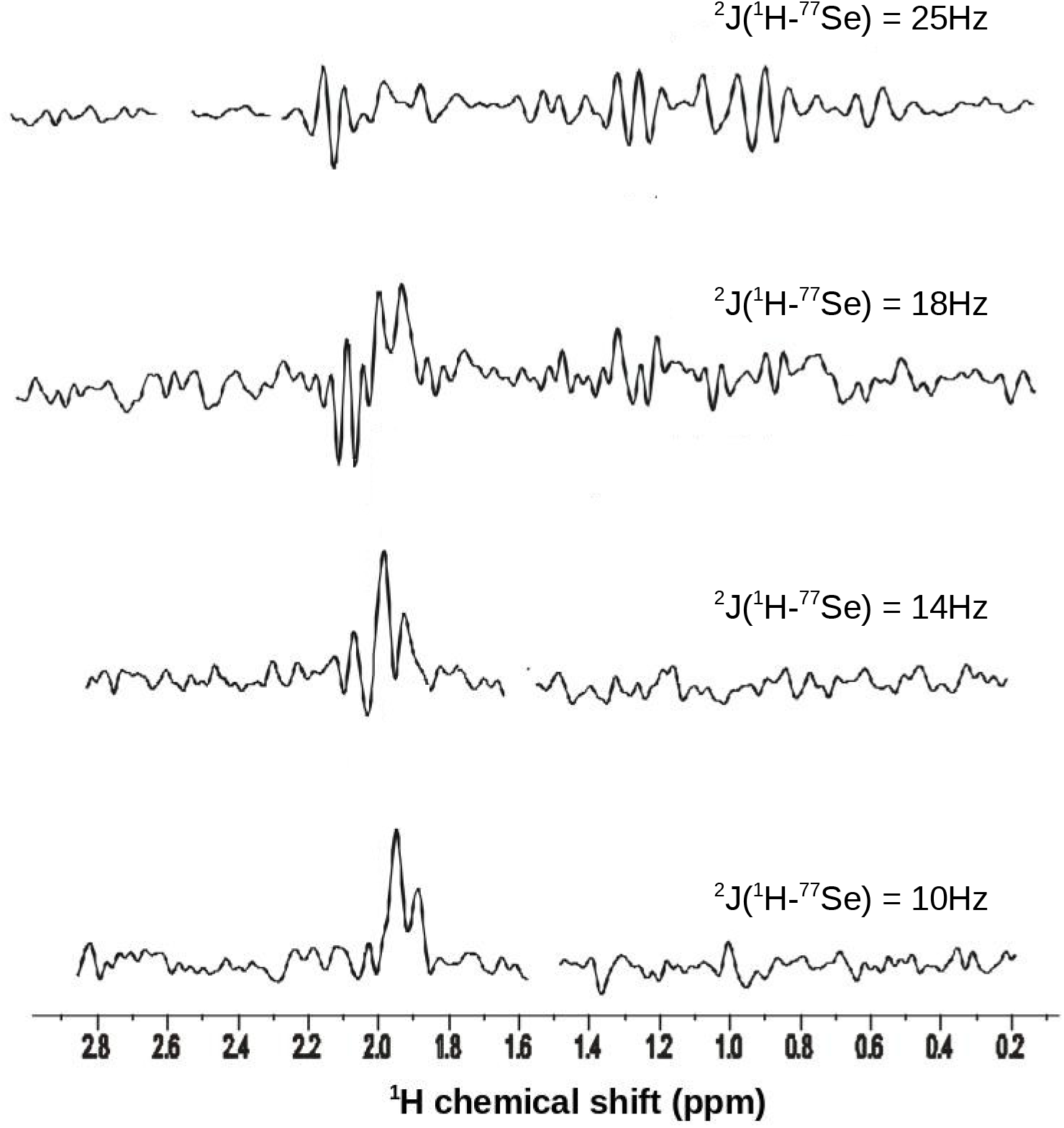
One-dimensional ^77^*Se* edited HMQC spectra of the reduced Se-Trx1 collected with the following ^2^ *J*(^1^*H* –^7 7^*Se*) coupling constants: 10, 14, 18 and 25 Hz respectively. The frequency of the carrier wave (O2P) was 150 ppm. The spectra were collected with 200 Se-Trx1 200μM, at 303 K temperature, with 2048 FIDs in a spectral window of 9615.38 Hz with 1024 points.

The 1D-HMQC spectra edited for ^77^*Se* of Se-Trx1, were collected with a greater number of FID accumulations, 3 lines of resonance between 2 and 1.6 ppm, were assigned as the *β*_1_ and *β*_2_ hydrogens of selenocysteines 30 and 33 from Se-Trx1 (figure 8).

**Figure 8.**
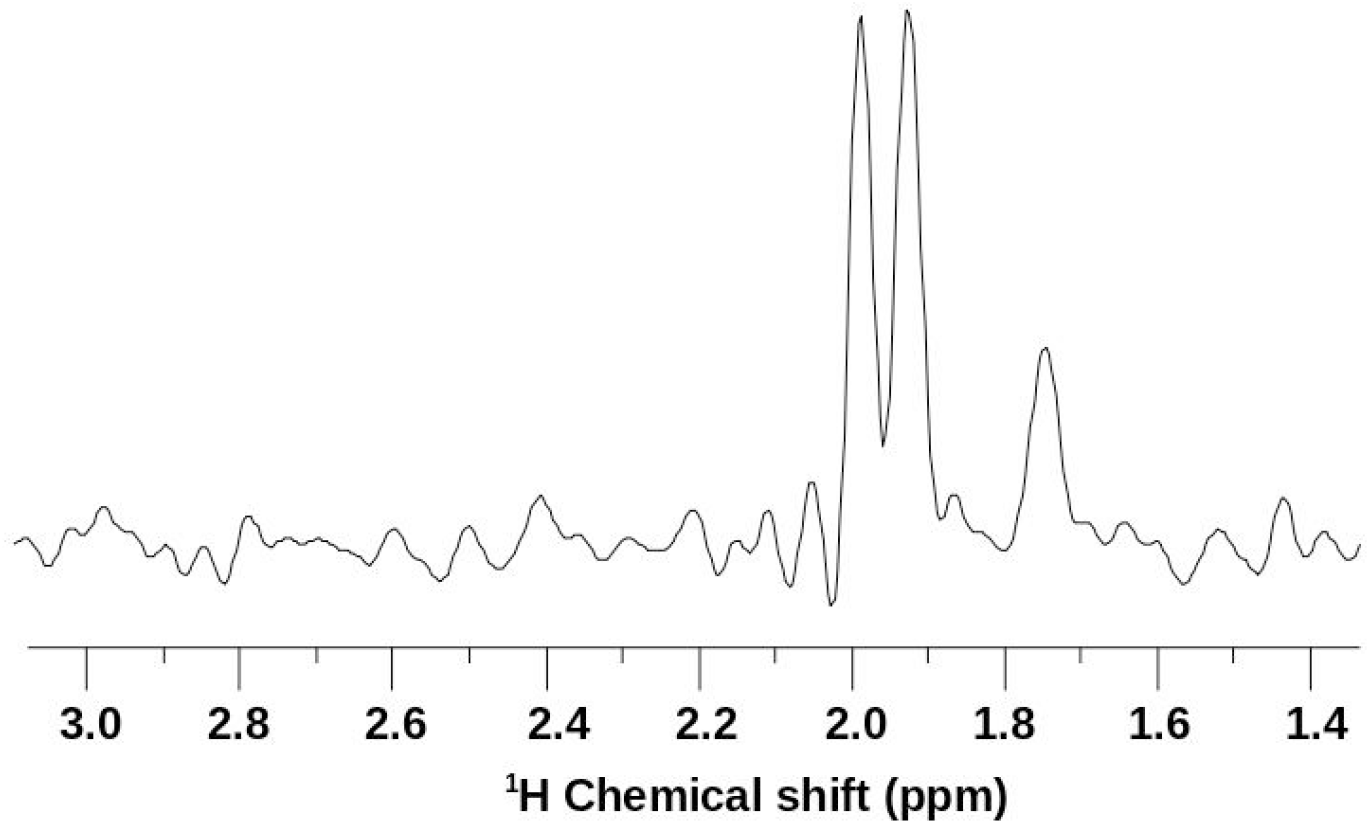
One-dimensional ^77^*Se* edited HMQC spectrum for 626μM Se-Trx1. The experiments were carried out at 303K temperature, using O2P=150 ppm and a coupling constant of 14Hz. A total of 20480 FIDs with 1024 points were collected, in a spectral window of 9615.38 Hz. The resonance lines observed between 2.0 and 1.6 ppm are probably generated by ^2^*J*(^1^*H* –^7 7^*Se*) couplings of the ^1^*H*(*β*_1_, *β*_2_) hydrogens of selenocysteines 30 and 33 from Se-Trx1

In order to assign the frequency of the ^77^*Se* signal coupled to the ^1^*H* protons in Selenocysteine as described above, we collected the two-dimensional HMQC (2D-HMQC) spectrum of Se-Trx1 as shown in the following figure (figure 9).

**Figure 9.**
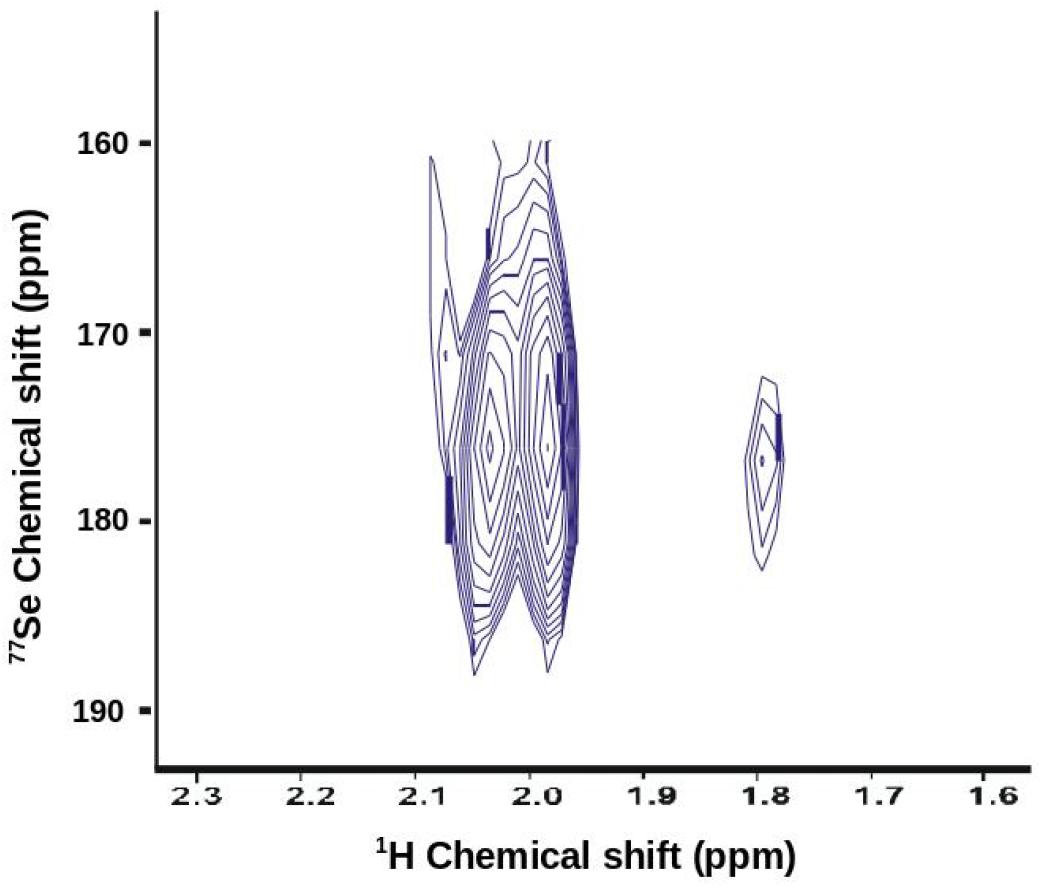
2D HMQC spectra edited to ^77^*Se* of the reduced Se-Trx1. The coupling between ^1^*H* and ^77^*Se* observed at approximately 175 ppm is probably from ^1^*Ĥ*_*β*1_ and ^1^*Ĥ*_*β*2_ and ^77^*Se* of the selenocysteines 30 and 33 from Se-Trx1

The two-dimensional HMQC spectrum for the ^1^*H*–^77^*Se* pair shows two ^1^*H* resonances between 2.1 and 1.9 ppm and a third resonance in approximately 1.8 ppm. These resonances are coupled with ^77^*Se* resonance at approximately 178 ppm.

#### Oxidized Se-Trx1

The 1D-HMQC ^1^*H* –^7 7^*Se* spectrum of the oxidized Trx1 was collected by varying the O2P as conducted with the reduced protein, however, the O2P variation used was between 150 and 1100 ppm. The modulation of the O2P in this frequency range was performed because previous works found ^77^*Se* resonances frequency in Selenosubtilisin under oxidizing and reducing conditions ranging between −220ppm in reduced protein and 1200ppm in oxidized protein [8]. We found a single resonance line at 2.6 ppm that probably encompasses the couplings ^1^*H*_77*Se*_ of selenocysteine 30 and 33 of Se-Trx1. This line was modulated by O2P frequency and presented the maximal intensity with O2P varying between 700 and 900 ppm (see figure 10).

**Figure 10.**
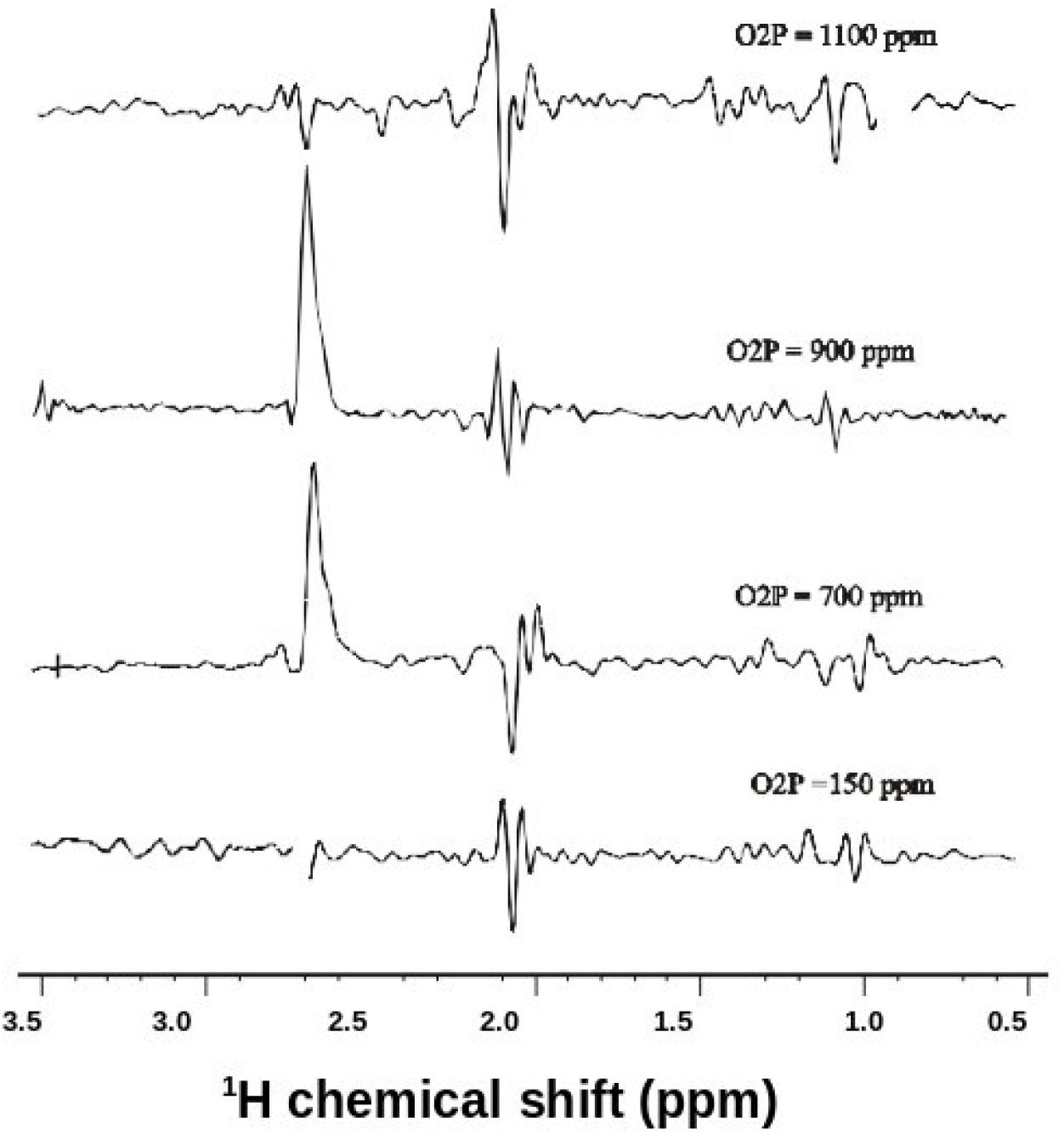
One-dimensional HMQC spectra edited to ^77^*Se* of the Se-Trx1 in 570 *μM* concentration. The Spectra were collected at 303K temperature by varying the O2P at 150, 700, 900, 1100 ppm respectively using a constant coupling of the 14Hz, 4096 FIDs were collected with 1024 points in a spectral window of the 9615,384 Hz.

The results found are in according with previous works that shows that the Oxidation of selenosubtilisin with hydrogen peroxide in the absence of thiol yields a species whose ^77^*Se* spectrum consists of two peaks at 1188 and 1190 ppm [8]. It shows that to different proteins, similar chemical environments provides ^77^*Se* chemical shifts in comparable scales.

In order to assign the ^77^*Se* resonance frequency coupled with ^1^*H*(*β*_1_, *β*_2_) hydrogens of selenocysteines 30 and 33 from oxidized Se-Trx1, we performed 2D-HMQC ^77^*Se* edited experiments; The 2D spectra shows ^77^*Se* resonances in approximately 830 ppm coupled with ^1^*H*_*β*_ protons in 2.3 ppm (see figure 11).

**Figure 11.**
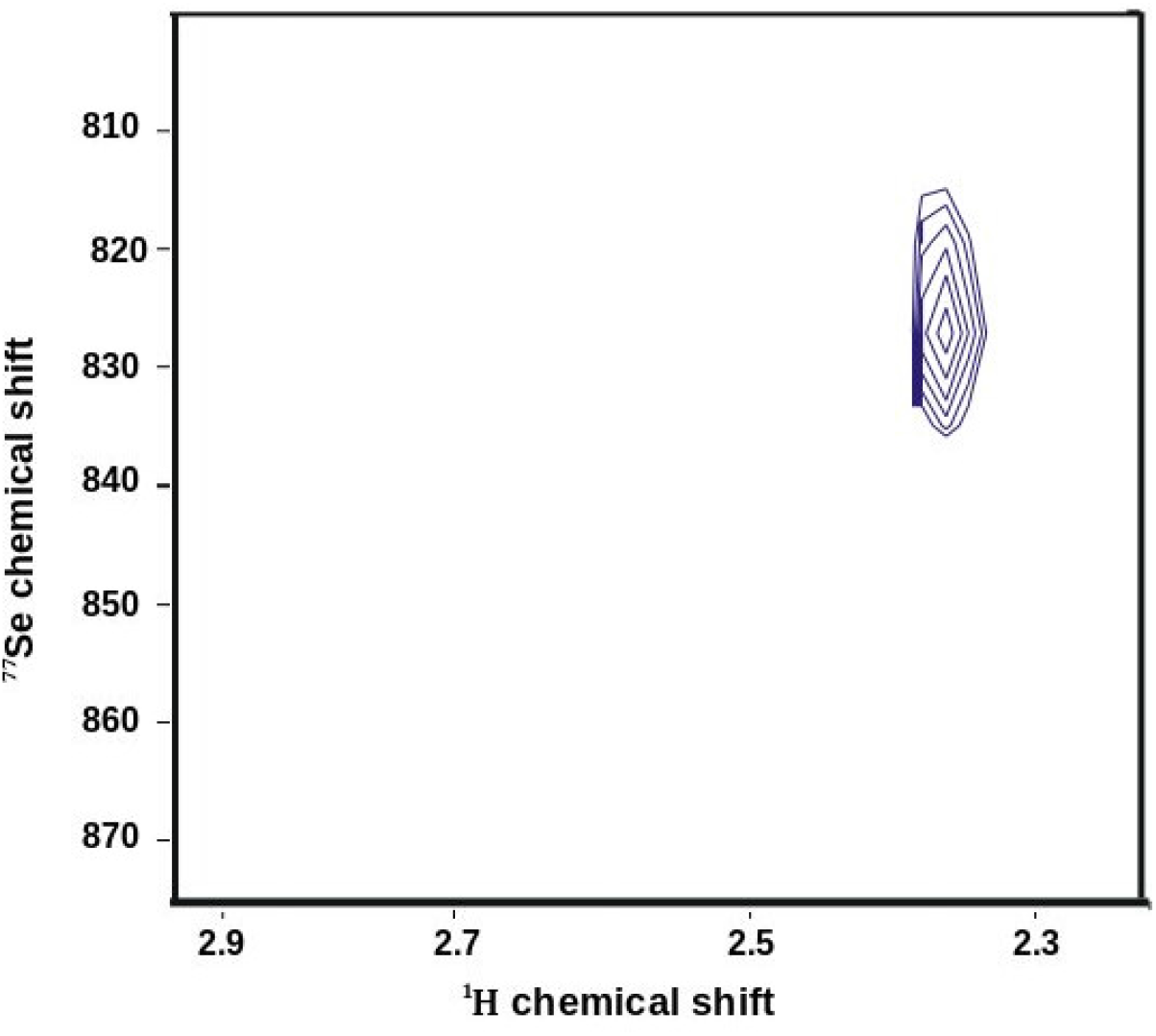
Two-dimensional HMQC spectra edited to ^77^*Se* of the oxidized Se-Trx1. The coupling found between 2.3 ppm and 830 ppm resonances can be assigned to ^1^*H*_*β*_ –^7 7^*Se* couplings in the Se-Trx1 diselenide bridge.

The large change in chemical shift found for ^77^*Se* resonances, between the oxidized and reduced forms of Se-Trx1 shows that the magnetic shielding of the ^77^*Se* atom in a protein is very sensitive to changes in the chemical environment. The disappearance of the separation of proton ^1^*H*_*β*1,*β*2_ resonances observed in the oxidized form if compared with the reduced form of Se-Trx1 is an indication that the oxidation generates the closure of the diselenide bridge, which was previously open with selenocysteines with their reduced selenol motifs. We suppose that the approximation of protons ^1^*H*_*β*1,*β*2_ to Selenium atoms generated by the closure of the diselenide bridge, determines the appearance of an broad resonance at approximately 2.6 ppm relative to coupling ^77^*Se* –^1^ *H_β_* in the Selenocistine of the oxidized Se-Trx1. The assignment of proton-coupled ^77^*Se* resonances through heteronuclear ^77^*Se* –^1^*H* HMQC experiments with the protein tagged with selenium isotopes even in natural abundance, as we did in this work, shows that this technique is extremely powerful to measure coupled ^77^*Se* –^1^*H* resonances in protein selenocysteines. Because ^1^_H_ –^77^*Se* coupling constants in the selenocystin dipeptide have already been measured, it is possible to access spin systems using coupling constants similar to those already measured in the selenocystin [18]. A demonstration of this fact is shown in the spectra of Figure 7 where we scanned coupling constants used in the pulse sequence of the heteronuclear HMQC and observed increases in the signal intensity when we used coupling constants close to those already measured in the spin systems of the l,l Selenocystin.

Although the assignment of ^1^*H* –^7 7^*Se* coupled resonances through heteronuclear NMR experiments have been done in 1,1 selenocystin spin systems, the same it not performed in whole proteins. It derives in the fact that a large chemical shielding response in ^77^*Se* nucleus brings about efficient relaxation routes, resulting in short transverse relaxation rates (*T*_2_) that broaden ^77^*Se* lines of the proteins in solution [6]. However, we show here that it is possible to obtain separate resonances for some ^1^*H* –^7 7^*Se* couplings in the selenocysteines from reduced Se-Trx1 (See Figures 7, 8, 9). The assignment of these resonances to protons ^1^*Hβ*_1,2_ coupled with the ^77^*Se* was made based on the coupling constants used in the heteronuclear HMQC experiments and those already measured in l,l Selenocystin. However, it does not seem possible the complete assignment of the spin systems from Selenocysteines of the Se-Trx1 with resonances totally separated by different coupling constants, due to the proximity between the values of the coupling constants and the broadening of the resonances generated by the chemical environment protein that is greatly affected by anisotropies. This problem is not pronounced for l-selenomethionine (Sem) whose chemical shift tensor span is *ω* = 580 ppm [16] and can be detected with ^77^*Se* at 7.5 percent natural abundance [28].

We detected, by mass spectrometry, different populations of Se-Trx1 whose Methionines were randomly substituted by Selenomethionines, so that between 0 and 4 of the Se-Trx1 Methionines were substituted with the Selenium atom (See Figures 1 and 4). It shows that the technique in biosubstitution used in this work could be extremely useful in the generation of different populations of Se-Trx1 for structural characterization using ^77^*Se* nucleus from different Methionines in several NMR strategies. It is possible that these different populations of Se-Trx1 can be purified to obtain specific structural information according to the position of Methionine residues replaced by Selenomethionines. Some works have used ^77^*Se* NMR to Probe different Protein environments of Selenomethionine [26]. In this experiments relevant conformational information is obtained combining ^77^*Se* Selenomethionine chemical shifts with x-ray cristalography data. The experiments shows that secondary and tertiary structures of all Selenomethionine (SeM) variants were not substantially impacted by the introduction of the SeM in different locations, and structural perturbations were restricted to SeM’s immediate vicinity. Although in this work we have not performed ^77^*Se* NMR experiments to evaluate the frequencies of Selenium resonances in selenomethionines, we have developed techniques that allow to obtain large amounts of information about the chemical environment of selenomethionines as it allows to obtain Se-Trx1 populations differentially labeled with Selenium relative to the position of Methionines; In the future, different ^77^*Se* NMR experiments could be conducted to obtain different structural information from Se-Trx1. This information can be used in the refinement of Trx1 structures obtained by NMR, in probing the dynamics of the enzyme from relaxation data of Selenium resonances at specific sites of Selenomethionines and also in studies of the interaction of this enzyme with molecular targets such as Ribonucleotide Reductase and Thioredoxin reductase for example; Finally, the results of this work allow to converge differential heterologous expression techniques of a protein with the objectives of specific structural information that one wants to obtain using ^77^*Se* NMR, this is therefore a powerful tool that can be used in probing relevant structural information using an isotope that presents an enormous spectral window subject to subtle interferences from magnetic shielding inside proteins.

## Acknowledgments

Coordenação de Aperfeiçoamento de Pessoal de Nível Superior (CAPES), Unidade Genômica do Instituito de Biofísica Carlos Chagas Filho-UFRJ, International Centre for Genetic Engineering and Biotecnology (ICGEB), Programas de Núcleos de Excelência (PRONEX) e Fundação de Amparo á Pesquisa do Estado do Rio de Janeiro(FAPERJ).

